# Emergence of multiple foraging strategies under competition

**DOI:** 10.1101/2024.04.19.590345

**Authors:** Hyunjoong Kim, Manoj Subedi, Krešimir Josić

## Abstract

Foraging strategies are shaped by interactions with the environment, and evolve under metabolic constraints. Optimal strategies for isolated and competing organisms have been studied extensively in the absence of evolution. Much less is understood about how metabolic constraints shape the evolution of an organism’s ability to detect food and move through its environment to find it. To address this question, we introduce a minimal agent-based model of the coevolution of two phenotypic attributes critical for successful foraging in crowded environments: movement speed and perceptual acuity. Under competition higher speed and acuity lead to better foraging success, but at higher metabolic cost. We derive the optimal foraging strategy for a single agent, and show that this strategy is no longer optimal for foragers in a group. We show that mutation and selection can lead to the coexistence of two strategies: A metabolically costly strategy with high acuity and velocity, and a metabolically cheap strategy. Generally, in evolving populations speed and acuity co-vary. Therefore, even under metabolic constraints, trade-offs between metabolically expensive traits are not guaranteed.

## 1. Introduction

Effective strategies for finding food and mates are central to survival and reproduction. Foraging strategies are determined by an organism’s phenotype, and evolve under environmental constraints including both intra- and interspecific competition [Stephens and Krebs, 1986]. Successful foraging requires perceptual acuity [Beauchamp, 2007; Kim and Lawley, 2024; Nummela et al., 2013] and the ability to navigate [Alexander, 2005; Herbert et al., 2020; Pontzer, 2016] the environment, both of which can be metabolically expensive [Laughlin et al., 1998; Niven and Laughlin, 2008; Wong-Riley, 2010; Hall et al., 2004; Pandolf et al., 1977; Ralston, 1958]. The optimal strategy for an isolated organism with full knowledge of the environment can be determined using the marginal value theorem [Stephens and Krebs, 1986; Parker and Smith, 1990]. This approach can be extended to competing organisms, but only when the birth and death rates for each are explicitly known [Chesson, 2000; Rocklin and Oster, 1976]. However, much less is known about how the environment and competition with other foragers shape the main phenotypic determinants of a foraging strategy: the ability to sense resources at a distance, and, the ability to move to the detected resource.

Animals occupying the same ecological niche use different foraging or hunting strategies with distinct metabolic demands. Even closely related organisms can adopt divergent strategies. Assassin bugs and ambush bugs have different hunting strategies, even though they are in the same family: While ambush bugs hide on flowers and lure their prey [Evans, 1931], assassin bugs hunt actively [Forthman and Weirauch, 2012]. Both hydra [Kaliszewicz, 2013] and limnomedusae [Colin et al., 2006] are in the same class, but hydra are sessile, whereas limnomedusae are motile and hunt actively. Such differences can be pronounced even between species occupying similar niches: Both sloths [Montgomery and Sunquist, 1975] and colobus monkeys [Chapman and Chapman, 2002] are herbivores and live in trees, but sloths move far more slowly, and have a lower metabolic rate [Nagy and Montgomery, 1980]. Thus, the same environment can lead to the evolution of distinct phenotypic attributes supporting foraging strategies with distinct metabolic demands. To explain the mechanisms that can lead to the emergence of such distinct foraging strategies here we introduce a minimal agent-based model of the coevolution of two important phenotypic determinants of foraging strategies: movement speed and perceptual acuity. Improving speed or acuity increases the rate at which resources are encountered, but also raises the metabolic rate. We assume that agents forage for food on a finite domain, and reproduce at a rate that depends on the resources they accumulated. We derive the optimal strategy for an isolated agent, and show this strategy is no longer optimal when organisms forage in groups. When we allow the agents to evolve and the influx of energy into the domain is constant, the surviving phenotypes have *lower* metabolic demands than an optimal, lone agent. Under these conditions, we also do not observe a trade-off between the two phenotypic attributes. Under certain conditions distinct phenotypes can coevolve with the population splitting into a smaller subpopulation using a metabolically costly foraging strategy with high speed and acuity, and a larger subpopulation using a metabolically inexpensive strategy with lower speed and acuity. We observe such coexistence when (i) the mutation rate is low so that phenotypic variance is sufficiently small for the two subpopulations to be distinguishable, and (ii) both resource density (per area) and energy density (per resource) are high enough so that both agents with low and high metabolic strategies can reproduce.

## 2 Methods

### 2.1 A stochastic agent-based model for the evolution of foraging strategies

Our results are based on the analysis, and stochastic simulations of a Markovian agent-based model that captures some of the fundamental processes that drive the evolution of foraging strategies [Gillespie, 1976; Champagnat et al., 2006, 2008; Doebeli et al., 2017; Liang and Brinkman, 2022]. The birth and death rates in our model depend indirectly on the success of the foraging strategy of an individual, and, through competition, on the foraging strategies of other individuals in the population. Successful foragers accumulate more resources, and are therefore more likely to reproduce and less likely to die. Offspring share phenotypic characteristics with their parents, and hence more successful foraging strategies are selected for.

We consider agents in a square domain, Ω, with periodic boundaries. Resources are randomly generated at points in the domain chosen with uniform probability in space, and temporal rate *λ* (figure 1). When consumed, each resource provides nutrients, described only in terms of metabolic energy. We label the agents in the domain at time *t* using indices *i* = 1, *· · ·*, *N* (*t*) where *N* (*t*) is the number of agents at time *t*. Each agent is characterized by a pair of phenotypic attributes, movement speed, *s*_*i*_, and perceptual acuity, *a*_*i*_. These attributes determine an agent’s phenotype, *ϕ*_*i*_ = (*s*_*i*_, *a*_*i*_), and hence its foraging strategy: Speed determines the rate at which an agent traverses the environment, while acuity determines the distance at which an agent can sense a resource. Hence, we will use the terms “phenotype” and “foraging strategy” interchangeably.

**Figure 1.**
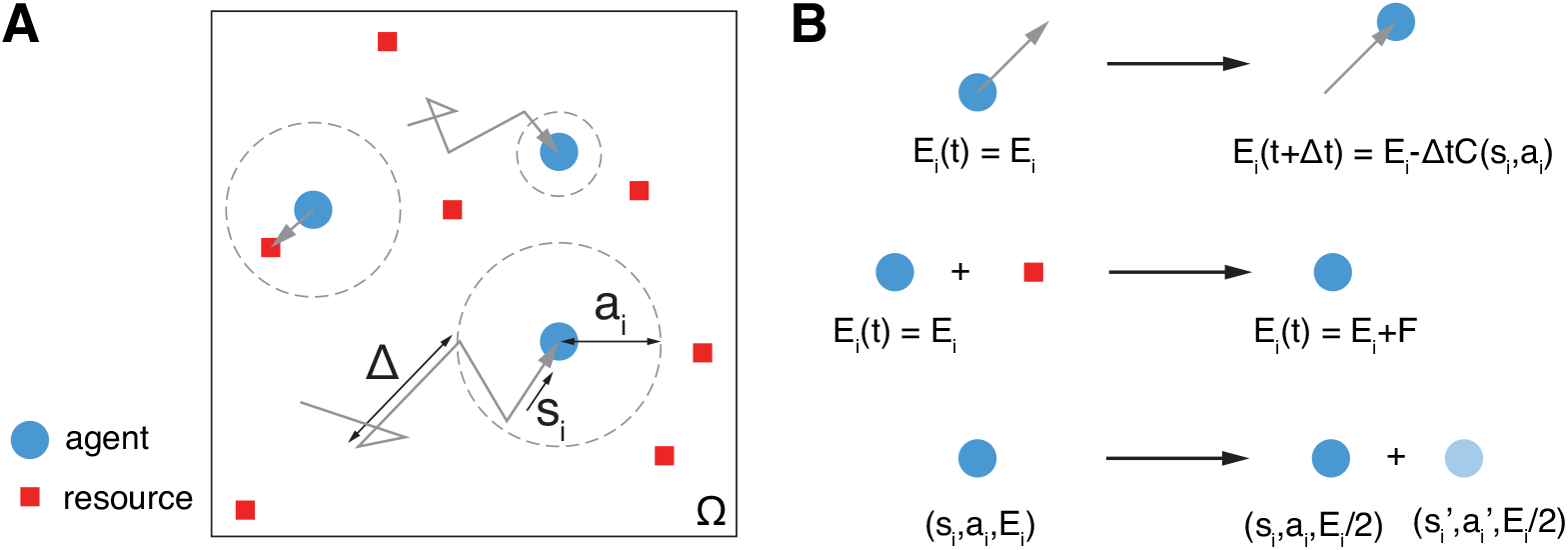
An agent-based model for the evolution of foraging strategies. (A) Agents forage on a rectangular domain, Ω, with periodic boundaries. Their foraging strategies are characterized by movement speeds, *s*_*i*_, and sensory acuities, *a*_*i*_. We assume that agents employ run-and-tumble motion. Distances between turns, Δ, are exponentially distributed with the same expected persistent length, *η*, so that Δ *∼* Exp(*η*). If an agent senses one or more resources, it moves ballistically towards the nearest resource, and fully consumes it once encountered. (B) An agent consumes energy at a constant rate depending on their foraging strategy (*top*). Consuming a resource adds an amount *F* to the agent’s stored metabolic energy (*middle*). Both birth and death rates are determined by the net energy the agent accumulated, *E*_*i*_ (*bottom*). Births produce one additional organism, with the phenotype of the offspring obtained by perturbing the phenotype of the parent. The parent and offspring split the parent’s stored metabolic energy equally.

We assume that agents employ *run-and-tumble motion* [Bressloff, 2021] with constant speed, *s*_*i*_. The times between changes in direction (tumbles), are exponentially distributed with a parameter, *f*_*i*_, that remains constant over the lifetime of the agent. All agents have the same expected persistent length *η* (the expected distance between turns). Hence, the tumbling rate depends on speed, *f*_*i*_ = *s*_*i*_/*η*. Upon reaching the end of a straight segment in its foraging trajectory an agent chooses a new direction by turning (tumbling) through an angle chosen independently and uniformly from [0, 2*π*). Thus, the spatial statistics of the foraging trajectories are identical for any speed, *s*_*i*_, and agents only differ in how fast they travel along these trajectories. If a resource is within a distance *a*_*i*_ of an agent, that is if an agent can sense a resource, then the agent moves ballistically toward the nearest resource with speed *s*_*i*_. Once agent *i* reaches a resource, an amount *F* is added to the agent’s stored metabolic energy, *E*_*i*_. All resources share the same energetic value, *F* .

Foraging for resources consumes energy at a rate that is determined by an agent’s phenotype, and hence by its foraging strategy. Agents always forage, and we only model the energy expended on foraging [Bennett and Gleeson, 1979; Stalmaster and Gessaman, 1984]. We assume that the metabolic energy of each agent, *E*_*i*_, is consumed at a rate given by

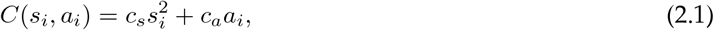

where *c*_*s*_, *c*_*a*_ are constants. The first term in equation (2.1) represent the rate of energy expenditure on movement which has been shown to be proportional to a power of speed [Alexander, 2005; Herbert et al., 2020; Pontzer, 2016]. Although the exact value of the exponent is not agreed upon, and likely depends on an animal’s size, mode of locomotion, and other factors, it is often assumed that movement consumes energy proportional to the square of the speed [Hall et al., 2004; Pandolf et al., 1977; Ralston, 1958; Subedi, 2023]. The second term in equation (2.1) represents the rate of metabolic expenditure of the sensory system, including the energy required to process sensory information [Beauchamp, 2007; Nummela et al., 2013]. We assume that this energy is proportional to the acuity, *a*_*i*_. In other words, an agent that can sense resources at twice the distance has a sensory system that expends energy at twice the rate.

The rates at which an agent reproduces and dies are determined by the available metabolic energy, *E*_*i*_. We use Hill functions to model the rate of asexual reproduction, and the death rate, respectively,

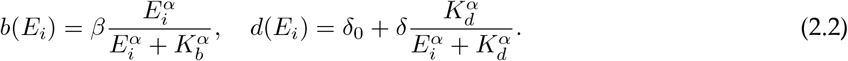

Here *K*_*b*_ (*K*_*d*_) determines the available metabolic energy at which the birth (death) rate is at half-maximum, *α* is the Hill coefficient, and *β* (*δ*) is the maximal birth (death) rate. In simulations we used *α* = 4. We also assume that agents die at a base rate *δ*_0_ regardless of how much energy they stored, and that *K*_*d*_ *< K*_*b*_. Hence, for a range of values of *E*_*i*_ an agent is likely to survive, but not likely to reproduce. Each birth results in one additional organism, and the parent and its offspring equally divide the parents’ stored energy. There is no additional loss of energy when an agent reproduces. A more realistic description of the metabolic rate in equation (2.1) may include other terms, which can be described by a constant offset. In turn, this could decrease birth rates and increase the death rates, and would be similar to the effects of increasing the base death rate, *δ*_0_, and shifting the half-maxima, *K*_*b*_ and *K*_*d*_.

To introduce variability in the phenotypes represented in the population, we either sample the phenotypic attributes at the outset of a simulation from some distribution, or let the offspring phenotype differ from that of their parent. In the second case, the speed and acuity of the offspring, (*s*_off_, *a*_off_), are given by a small perturbation of the parent’s phenotype, (*s*_par_, *a*_par_):

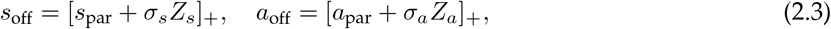

where *Z*_*s*_ and *Z*_*a*_ follow the standard normal distribution. The parameters *σ*_*s*_ and *σ*_*a*_ describe the variability of the two attributes across generations [Hayman and Mather, 1955; Mather and Jinks, 1971]. Square brackets denote rectification, [*x*]_+_ = *x* for *x ≥* 0 and [*x*]_+_ = 0 for *x <* 0, ensuring that phenotypic attributes are non-negative. Agents that do not move, *s* = 0, or cannot sense, *a* = 0, are not able to forage, and die without reproducing. In addition, we assume that offspring start at a location, (*x*_off_, *y*_off_), at some distance from their birthplace, (*x*_par_, *y*_par_):

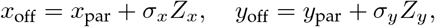

where *Z*_*x*_ and *Z*_*y*_ follow a standard normal distribution and *σ*_*x*_ and *σ*_*y*_ are small constants. This perturbation breaks up spatially synchronous clusters that form when offspring and their parents occupy the same initial point in space and move towards the same resources in high resource density environments. We implemented this model in Matlab [Kim and Subedi, 2024], and the code is openly available.

## 3 Results

We use an agent-based model to address key questions about the co-evolution of phenotypic attributes and the resulting foraging strategies: How do characteristics of the environment shape foraging strategies with different metabolic demands? Can populations with different strategies coexist in the same environment or niche? To address these questions we model agents foraging for calorically rich resources in a finite domain. The agents’ rates of reproduction and death depend on the amount of resources (energy) they are able to accumulate. Their foraging strategies and metabolic rates are determined by two phenotypic attributes, speed and acuity, both of which evolve under environmental pressures, as well as competition with other agents. We first describe the optimal phenotype for a single forager. We next ask what distributions of phenotypes are most successful in phenotypically homogeneous population (§3.2), and in phenotypically heterogeneous populations, where all agents expend energy at an equal rate on foraging (§3.3), and, finally, without constraints on metabolic cost (§3.4).

### 3.1 Optimal foraging of individual agent

Determining which strategies maximize the rate at which energy is accumulated is a core question of optimal foraging theory. The problem is typically formulated in terms of the *global capture rate* [Stephens and Krebs, 1986], defined as the net energy gain determined by the rate at which resources are gathered and the rate at which energy is expended. To find the strategy that maximizes the global capture rate for a single agent, we numerically compute the resource uptake rate, determining the rate at which an agent gains energy, and then examine the asymptotic behavior of the resource uptake rate at low and high acuity. We then analytically characterize the maxima of the global capture rate. We will show subsequently that the optimal strategy for an isolated agent is no longer optimal when foraging in a group.

We start by determining the average rate at which a single agent encounters resources, *ν*(*s, a*|*r*), as illustrated in figure 2A. This rate depends on the agent’s speed, *s*, and acuity, *a*, as well as the number of available resources, *r*, in the domain. When resources are generated at a constant rate, the number of available resources fluctuates around an equilibrium value^1^, *r*^*∗*^, at which the agent consumes resources at the rate at which they appear, so that *ν*(*s, a*|*r*^*∗*^) = *λ*. To simplify the analysis we therefore consider an environment with a constant number of available resources, *r*, although our conclusions hold more generally. Since we assume a fixed persistent length, *η*, the spatial statistics of the search trajectories are identical for any *s*. This implies that the encounter rate takes the form

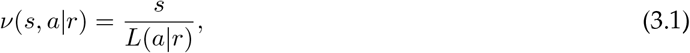

where *L*(*a*|*r*) is the average length of a trajectory between target encounters when the agent’s acuity is *a* and there are *r* resources in the domain. Hence, the resource encounter rate is linear in speed. The average distance between an agent at location **x**_0_ and *the nearest* of *r* uniformly distributed resources in the periodic domain, Ω, is

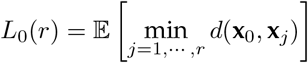

where **x**_*j*_ *∼ U* [0, 1]^2^ for *j* = 0, *· · ·*, *r* and *d*(**x, y**) is the shortest (Euclidean) distance between **x, y** in the finite periodic domain [0, 1]^2^. Here **x**_*j*_ is the location of resource *j*. By assumption, an agent can detect a resource within a radius *a*.

**Figure 2.**
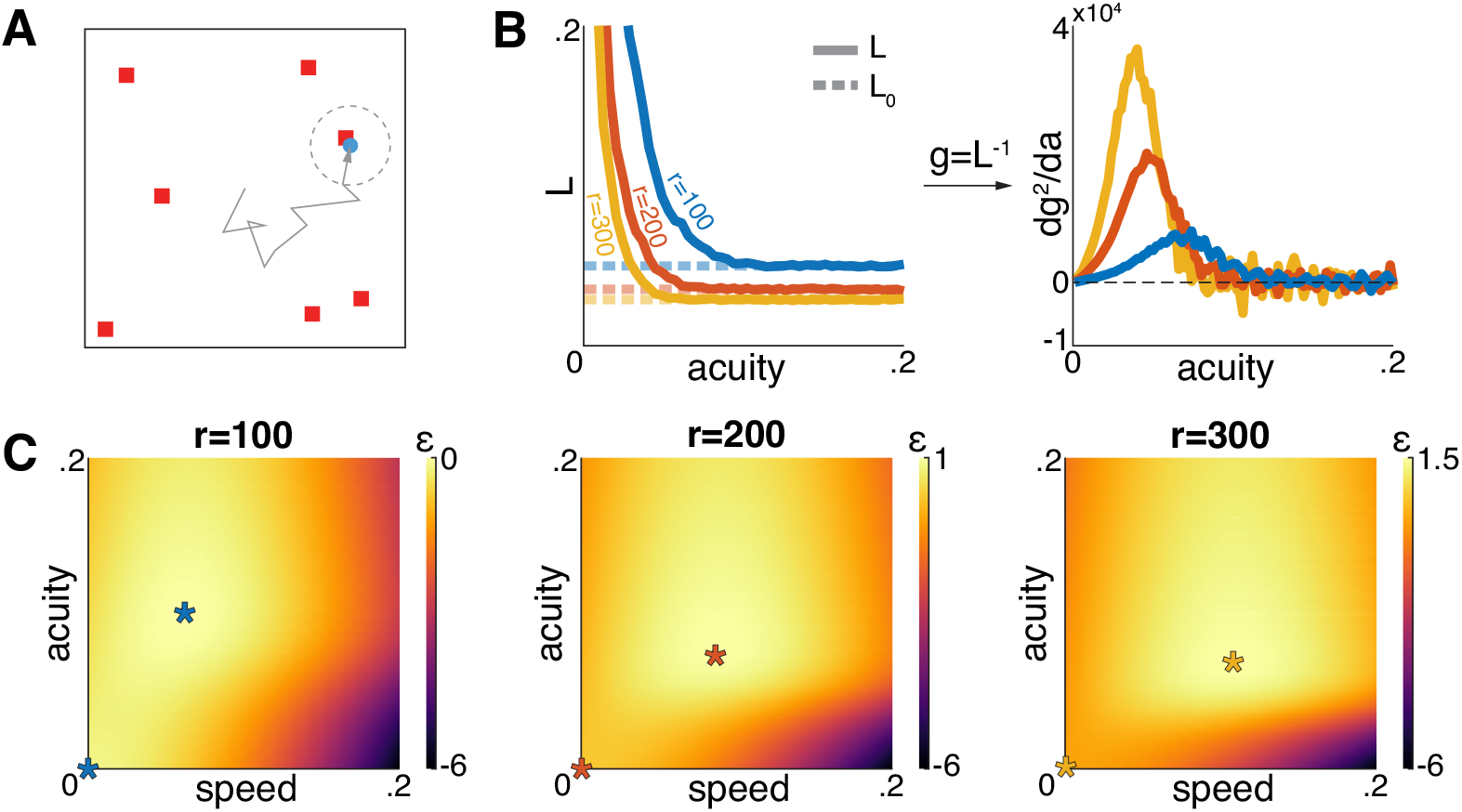
Optimal foraging strategy of a single agent. (A) A single agent forages for a fixed number, *r*, of resources in the domain. The net energy gain of the agent depends on the average length of the trajectory between resource encounters, *L*. (B) As acuity increases, *L* converges to the average minimum distance, *L*_0_, defined in equation (3.2), and movement between resource locations becomes ballistic (*left*). Conversely, decreasing *a* results in random run-and-tumble motion and search trajectories of increasing length, with *L* diverging to infinity (see the supplementary material). The shape of *L*(*a*) determines the existence of an optimal foraging strategy by Theorem 1 (*right*). (C) Two local maxima of the net energy gain, one at *s* = *a* = 0 and the other defining the optimal foraging strategy. Parameters are listed in appendix.

If *a ⪢ L*_0_(*r*), the agent always senses a resource, and does not engage in random search. Therefore, for large *a* we expect

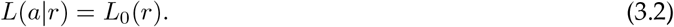

This agrees with numerical simulations (figure 2B). The resource encounter rate thus increases as acuity increases, and saturates when acuity is sufficiently high for an agent to always sense a resource.

If *a ⪡ L*_0_(*r*), an agent placed randomly within the domain is unlikely to sense a resource. A simple argument described in the supplementary material shows that *L*(*a*|*r*) diverges as *a* approaches zero at the same rate or faster than *a*^*−*1^.

The global capture rate, *i*.*e*., the average net energy gain of the agent can be defined by

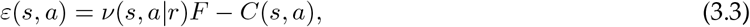

where the first term represents the energy gain rate of the agent, while the second is the metabolic energy expenditure rate for an agent with phenotypic attributes *s* and *a*.

Simulations show that, given a fixed number of resources, *r*, there are two local maxima for the global capture rate (figure 2C). We can prove this result under reasonable assumptions (see proof in the supplementary material):

#### Theorem 1

*For a fixed r, ε*(*s, a*) *has two local maxima in* (*s, a*) *∈* [0, *∞*)^2^, *one in the interior of the phenotype space, ϕ*_*opt*_ *∈* (0, *∞*)^2^, *and the other at the origin, ϕ*_0_ = (0, 0), *if g*(*a*) := 1/*L*(*a*|*r*) *has a unique solution a*_*opt*_ *such that*

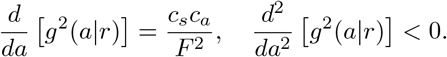

We verified numerically that *g*(*a*) satisfies the condition of this theorem (figure 2B) when *c*_*s*_*c*_*a*_/*F* ^2^ is sufficiently small. Since *ε*(*ϕ*_0_) = 0 and *ε*(*ϕ*_opt_) *>* 0, *ϕ*_opt_ is the optimal phenotype for an individual forager.

At equilibrium, the amount of resources in the domain is approximately constant, and *ϕ*_opt_ remains optimal if resources are supplied at a constant rate. However, when sufficiently many agents forage in the domain they start competing for resources, and *ϕ*_opt_ may no longer result in the highest capture rate.

### 3.2 Steady-state of a population using single strategy

Does the optimal strategy for a single forager maximize the size of a phenotypically homogeneous population? A metabolically costly phenotype could lead to a higher global capture rate. Conversely, metabolically inexpensive phenotypes could allow for a large population even with a low capture rate. We next determine the steady-state population size for a given *fixed* phenotype by introducing a mean-field energy balance equation. This equation helps explain why optimal single forager phenotypes do not maximize the size of phenotypically homogeneous populations.

To determine the size of the steady-state population of identical agents with phenotype *ϕ*, we balance the energy influx to the environment with the energy consumed by the agents. We assume the population is in equilibrium and denote the energy accumulated by each agent averaged over time and across the population by *E*^*∗*^, the population size by *n*^*∗*^, and the number of resources by *r*^*∗*^. We assume each agent has an equal amount of net energy at their disposal equal to the population average, *E*_*i*_ = *E*^*∗*^.

Agents must balance the rate of energy accumulated while foraging with the energy rate expended on foraging, *C*(*ϕ*), and reproduction

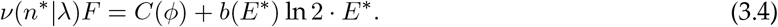

Here *ν*(*n*|*λ*) is the resource encounter rate of a single agent when *n* agent forage in a domain with resources generated at rate *λ*. The second term on the right-hand corresponds to the loss at reproduction since accumulated energy is divided equally between parent and offspring at birth. At equilibrium, birth and death rates are equal,

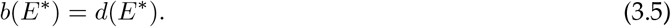

This is an implicit equation which determines the accumulated energy at equilibrium, *E*^*∗*^. Lastly, the resource generation rate and the total resource encounter rate by all agents are also balanced at equilibrium,

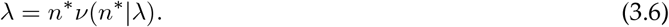

Substituting (3.6) into (3.4) gives the steady-state population size,

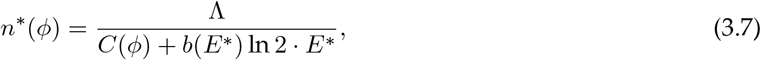

where Λ *≡ λF* represents the total energy influx to the system. The denominator represents the average net energy rate expended by an individual agent. Thus, equation (3.7) determines the expected number of agents in a domain given that each agent has at its disposal an amount of stored energy *E*^*∗*^ given implicitly by equation (3.5). We show in the supplementary material how to estimate the amount of resources available to the agents. Equation (3.7) also provides a bound on the phenotypes with high metabolic costs that an environment can support since *n*^*∗*^(*ϕ*) *⪡* 1 if *C*(*ϕ*) *⪢* Λ.

Surprisingly, in homogeneous populations competition for resources does not impact population size: Equation (3.7) shows how the steady-state population of identical agents depends on the metabolic rate when resources are generated at rate *λ*. Equation (3.5) shows that the accumulated energy at equilibrium, *E*^*∗*^, does not depend on the environment or the population phenotype. Therefore, equation (3.7) implies that the steady-state population, *n*^*∗*^, depends only on the phenotype’s metabolic expenditure rate, *C*(*ϕ*), and the energy influx into the system, Λ. Thus, two homogeneous populations with different phenotypes, *ϕ*_1_ and *ϕ*_2_, but equal metabolic rates, *C*(*ϕ*_1_) = *C*(*ϕ*_2_), will reach approximately equal population sizes at equilibrium. These observations agree well with numerical simulations (figure 3B).

**Figure 3.**
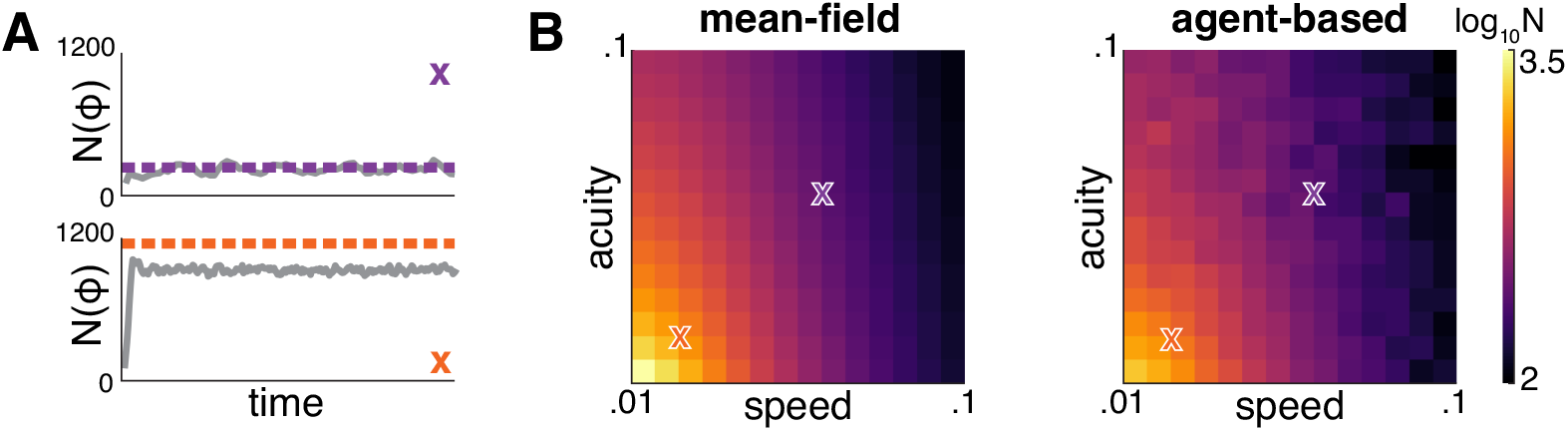
Steady-state size of a homogeneous population. (A) Stochastic simulations (*solid curves*) and the mean-field approximation in equation (3.7) (*dashed lines*) agree well for two example phenotypes with high (*top*) and low metabolisms (*bottom*). The mean-field approximation overestimates the steady-state population size because it does not account for random deaths of agents with low accumulated energy. (B) The mean-field approximation (*left*) predicts the equilibrium population in agent-based simulations (*right*) well in homogeneous populations showing how population size depends on phenotype. The two example phenotypes in (A) are denoted by crosses. Parameters are listed in appendix.

With fixed phenotype the steady state population size increases with a decrease in metabolic rate (figure 3B) since *n*^*∗*^(*ϕ*) depends inversely on the metabolic cost *C*(*ϕ*). Therefore, the phenotype that results in the largest homogeneous population is not the optimal phenotype for an individual agent discussed in the previous section.

Does this imply that multiple phenotypes (foraging strategies) with equal metabolic cost can coexist? Do phenotypes with low metabolic costs outcompete more costly phenotypes? The answers depend on how competition impacts foraging rate in heterogeneous populations. We next show that one phenotype outcompetes others with the same metabolic costs.

### 3.3 Selection between strategies with the same metabolic cost

In heterogeneous populations different phenotypes compete for resources. However, it is unclear whether under such competition a single, dominant strategy displaces others, or if multiple strategies can coexist.

To address this question, we first consider competition between agents using foraging strategies with equal metabolic costs. To simplify our analysis we assume that parents give birth to phenotypically identical offspring. We ask which of the phenotype or phenotypes initially present in the domain survive in the long term. Expressions for the resource encounter rate are difficult to obtain, and we therefore proceed with numerical simulations.

We first determine the steady-state population distribution over the multiple strategies as we did in §3.2. Again, we consider the mean-field approximation for a heterogeneous population at equilibrium. Let *ϕ*_*i*_ for *i* = 1, 2, *· · ·* denote phenotypes with the same metabolic rate, *C*. We assume that all agents that share the same phenotype also have equal average accumulated energy and resource encounter rates. The rate at which energy is accumulated for agents with phenotype *ϕ*_*i*_ must again balance the rate at which energy is expended on foraging and reproduction:

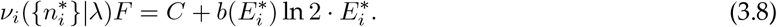

Here *ν*_*i*_({*n*_*i*_*}*|*λ*) is the resource encounter rate for an agent with phenotype *ϕ*_*i*_, *i* = 1, 2, *· · ·* under competition with *n*_*i*_ subpopulations with phenotype *i* = 1, 2, *· · ·*, and 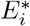 is the average energy accumulated by this phenotype. At equilibrium,

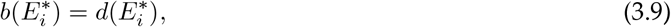

for all *i*. Equation (3.9) has a unique solution, *E*^*∗*^, and so *E*_*i*_ *≡ E*^*∗*^ for all *i*. Balancing the resource generation and consumption gives

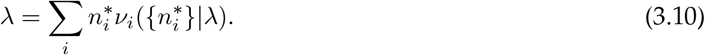

Substituting (3.8) into (3.10) gives the total steady-state population

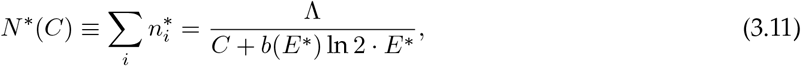

which has the same form as equation (3.7). The population size, *N* ^*∗*^, is determined by the metabolic expenditure rate of agents. However, the subpopulation sizes, 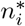, are determined by the resource encounter rates, *ν*_*i*_. These rates are difficult to obtain analytically, and we therefore resort to numerical simulations.

Agents at equilibrium have intermediate speed and acuity, when the initial populations consist of diverse phenotypes with equal metabolic cost, as illustrated in figure 4A. We started our simulations with a population of *∼*10^3^ agents with phenotypes sampled from contours defined by constant metabolic rate, *C*, in phenotype space. The absence of mutations implies that the population is confined to the same contour for all time. Figure 4C shows that in the long term the surviving phenotypes cluster around a single, “dominant” strategy with intermediate speed and acuity. However, at high metabolic rates, *C*, the population goes extinct after a short time (figure 3C).

**Figure 4.**
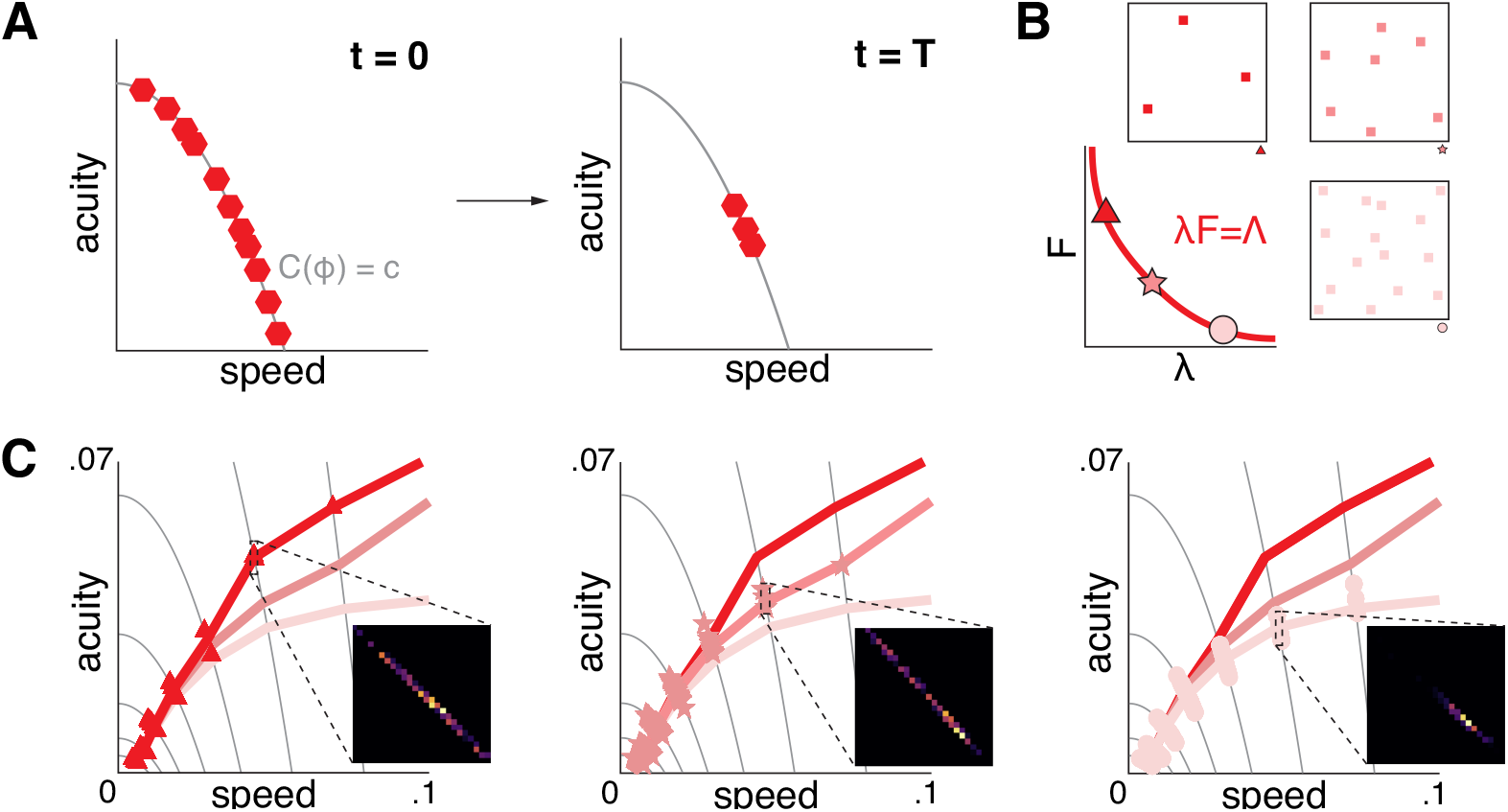
Competition between foraging strategies with equal metabolic cost. (A) Simulations indicate that a tight cluster of “dominant” foraging strategies outcompetes other strategies with equal metabolic cost. These strategies are characterized by intermediate speed and acuity. (B) Three different resource environments with the same energy influx. (C) Population averages of the phenotypes of the surviving strategies with different initial metabolic costs (*solid curves* connect strategies with equal metabolic cost) in the three different environments: High *λ* and low *F* (*left*), intermediate (*middle*), and low *λ* and high *F* (*right*). The inset panels show the phenotype distributions of the surviving strategies pooled over different simulations. High speed and acuity are selected for when resources are scarce. Parameters are listed in appendix.

The observation that a balance between speed and acuity is best for foraging is intuitive as agents with intermediate phenotypes have higher resource encounter rates than those with more extreme phenotypes. An agent with high speed and low acuity wanders randomly, and may take a long time to detect a resource. A slow moving agent with high acuity can detect distant resources, but may not reach them before they are scooped up by faster moving agents.

We next asked how environmental factors affect the dominant strategy. To address this questions we varied the resource generation rate, *λ*, and the energy delivered by a resource, *F* . We set *λF ≡* Λ so that the total energy influx to the domain was constant, as illustrated in figure 4B. When resource density is high the dominant phenotypes are characterized by high speed and low acuity. In contrast, a low density of energy rich resources favors high acuity and lower speed (figure 4C). When resources are abundant, but have low energy density, agents do not require high acuity to detect them. However, high speed allows the agents to collect the resources at a higher rate. When resources are sparse, on the other hand, acuity becomes more important.

Thus, simulations indicate that multiple strategies with the same metabolic cost do not co-exist at equilibrium. Under competition, a single strategy leads to the highest resource encounter rate, and outcompetes all others. We next ask whether this observation also holds in evolving populations.

### 3.4 Coexistence of phenotypic attributes

We next assume that there is a small difference in phenotype between parent and offspring modeling the effect of mutations. We use simulations to characterize the distribution of phenotypes at equilibrium, and show that speed and accuracy are often correlated, and the population can split into two subgroups with distinct phenotypes. We again consider environments with different resources densities (figure 4B). To do so we fix the energy influx into the environment, *λF* = Λ, and vary *λ* and *F* = Λ/*λ*. We use Monte Carlo simulations to approximate the long term behavior of the population (See figure 5A and Appendix for more details).

**Figure 5.**
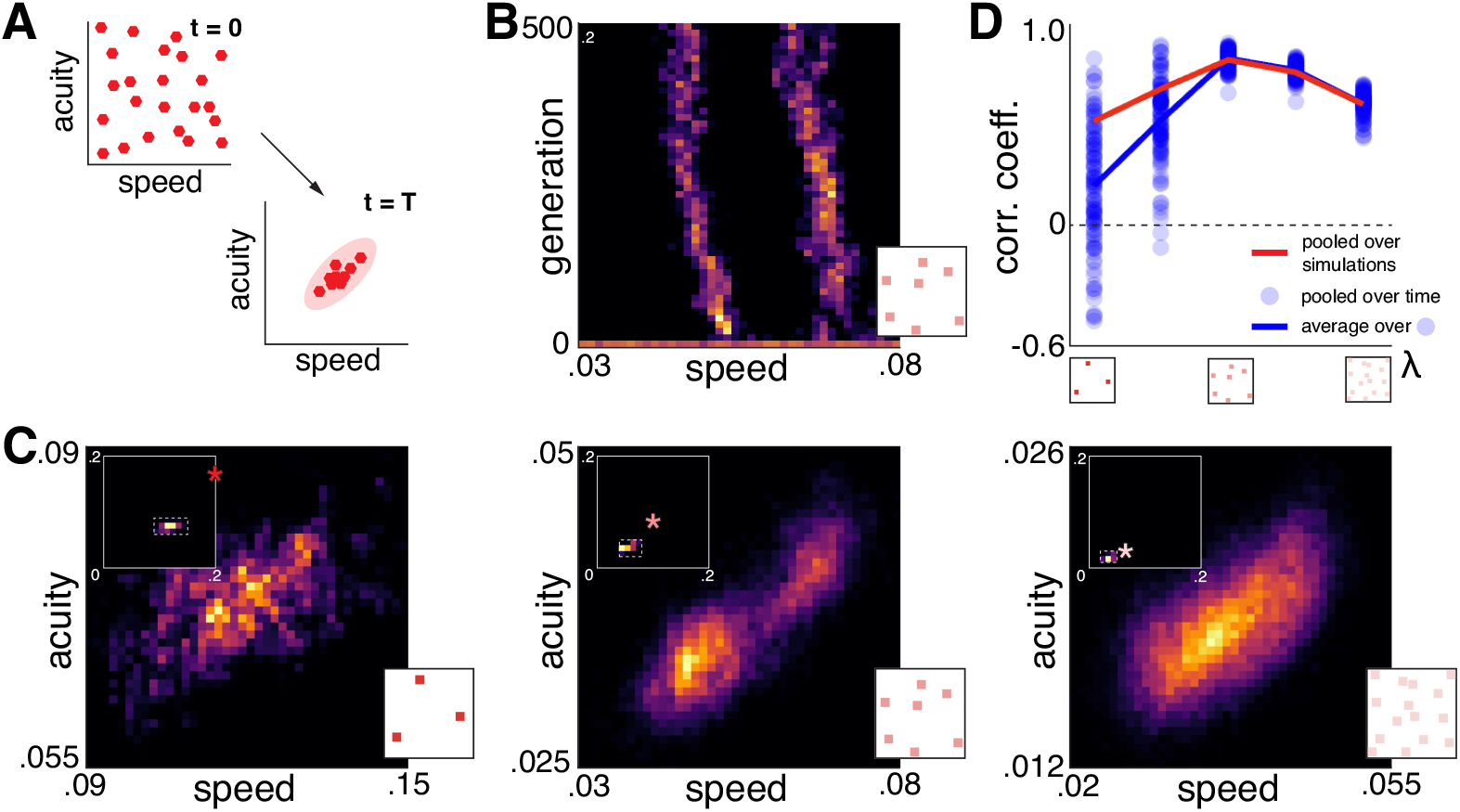
The equilibrium distribution of phenotypes depends on resource density. (A) Without constraints on metabolic costs, selection results in convergence to a quasi-stationary distribution. (B) A realization showing that at intermediate resource density two distinct phenotypes coexist over many generations (not shown are the acuities which also remain distinct between the groups). (C) Equilibrium distributions of phenotypic attributes pooled over simulations in different environments. Low *λ* and high *F* environment (*left*) and the high *λ* and low *F* environment (*right*) support a unimodal distribution of phenotypes, while intermediate environments (*middle*) lead to a bimodal distribution. These equilibrium distributions correspond to strategies that are metabolically cheaper than the optimal strategy of a single forager in the same environment (*stars in the subpanels*). (D) Correlation coefficients between the two phenotypic attributes at equilibrium. To compute the correlation coefficients, we pool phenotypes over simulations at a fixed time (*red curve*), and over time in individual simulations (*blue circles*), and average over simulations and time (*blue curve*). Pooling over time results in lower correlation coefficients when resource density is low. Parameters are listed in appendix.

Without a constraint on metabolic costs, we again observe that resource-sparse environments favor metabolically costly phenotypes. Moreover, different foraging strategies can co-exist at equilibrium in moderate environments (figure 5B and supplementary material, figures S1 and S2). Here, the population converges to two clusters composed of phenotypes with different metabolic costs. When resource density is low or high we observe a single cluster in phenotype space. At high resource densities, energetically cheap strategies have an advantage. When resources are sparse, but calorically rich, metabolically costly foraging strategies dominate.

The strategies that are selected for under competition and mutation are *less metabolically costly* than the optimal strategies of isolated foragers in the same environment (subpanels in figure 5C). For this comparison, we used the steady-state quantity of resources in the crowded environment, and obtained the optimal single agent strategy as described in §3.2. The ratio between the metabolic cost of the optimal individual strategy, *ϕ*_opt_, and the average metabolic cost of the population in the crowded environment, 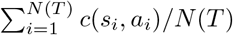, is approximately 2.8, 3.0, and 2.7, in the order of panels in figure 5C, respectively. Since metabolically costly phenotypes survive for shorter periods when they fail to encounter resources, failure to find resources due to competition impacts such phenotypes more strongly.

We also find that speed and acuity tend to be positively correlated at equilibrium. A negative correlation would correspond to a trade-off between the two phenotypic attributes. In §3.3 our assumption of fixed metabolic cost forced such a trade-off. Without this constraint, at equilibrium the population consists of phenotypes with positively correlated attributes (figure 5D), particularly at moderate resource densities.

At low resource densities convergence to an equilibrium distribution is very slow, and pooling over simulations at a fixed time gives different results than pooling over many snapshots from single realization. In particular, the correlation coefficient between the attributes depends on if we pool the phenotypes, *ϕ*_*j*_(*t*), at some time *t ⪢* 0 across simulations *j*, to obtain 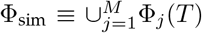, or across time,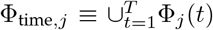 . The average correlation coefficient when pooling over time and simulations, 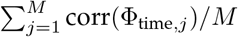, is lower than the correlation coefficient when pooling over simulations, corr(*ϕ*_sim_) (figure 5D). This indicates that each realization is frozen for a long time in a state where the two phenotypic attributes are weakly correlated. However, these frozen states are not the same across simulations, but are centered approximately along a line with positive slope in phenotype space. We expect this agent-based system is ergodic, and hence this difference may reflect very slow mixing when resource densities and population sizes are low (supplementary material, figure S4). This difference does not occur at intermediate and high resource densities where equilibrium populations are larger (figures 5D and supplementary material, figure S3).

The variability in the offspring phenotype has a large impact on the variance of the phenotype distribution at equilibrium. If the variability is too large the equilibrium distribution ceases to be bimodal, as the two modes are hard to distinguish (supplementary material, figure S5). If the variability is very low and the initial phenotypes are chosen to be close to one peak of the equilibrium distribution, then it can take long time to reach the other peak.

## 4 Discussion

We introduced an agent-based model of foragers competing for limited resources on a finite domain. The foraging strategies of the agents in the model are determined by two abstract phenotypic attributes: the ability to sense resources (acuity), and the speed at which the agents traverse the environment. The two attributes are complementary: While acuity helps in locating and thus reaching resources efficiently, speed is essential for searching, and outracing other foragers who may have spotted the same resource. The two attributes are thus both essential for foraging and successful competition with other agents.

We assumed that increased speed and acuity result in higher energetic expenditures [Laughlin et al., 1998], so that the resulting metabolic demands shape the evolution of these attributes. The resulting model is computationally and semi-analytically tractable and allowed us to address several fundamental questions: How are limited metabolic resources divided to support both sensory processing and movement? Are there trade-offs between the demands of these two systems, or do they co-evolve? Which environments support the evolution of large populations of organisms with low metabolic demands, and which environments support metabolically costly foraging strategies? Can two distinct strategies co-exist? We found that speed and acuity tend to be correlated in a population. This is not surprising, as speed and acuity complement each other in foraging and competition: An agent who can see farther can afford to move quickly, as it often moves towards a resource. If the flux of energy is fixed, the environment supports few metabolically expensive agents with high acuity and speed, many energetically inexpensive agents with low acuity and speed, or a range of phenotypes interpolating between these extremes. Which of these phenotypes has an advantage depends on how difficult it is to find resources, with abundant, but calorie deficient resources favoring large populations of slow moving, short-sighted agents. Strategies that emerge under competition are less metabolically costly than the optimal strategy of a lone forager. This is in contrast to previous findings that competition between individuals has little impact on human brain size, and, presumably, on the ability to process sensory information [González-Forero et al., 2017; González-Forero and Gardner, 2018].

Surprisingly, multiple phenotypes can coexist at equilibrium in homogeneous and patchy resource environments (results not shown for patchy environments, but see [Subedi, 2023]). This is similar to the bimodality in the distribution of ornament sizes [Clifton et al., 2016], which can emerge because it may be more efficient to adopt a metabolically less expensive phenotype than trying to compete directly with more costly phenotypes. This suggests that our model could also be extended to multiple species evolving in environments where different types of resources are available.

Although the full model is not analytically tractable, we were able to analyze several of its aspects: The carrying capacity of the environment can be obtained directly as function of the available resources, and the metabolic demands of a phenotype. Thus, our abstract formulation paid dividends, and further analysis of the model may be possible. An approximate mean field model was suggested in [Subedi, 2023]. Moreover, phenotypes evolve on a timescale that is far longer than the timescale of foraging dynamics, and this separation of timescales may offer another way to analyze the system. The main difficulty in such analysis is accurately describing competition between agents, which will be challenging using standard mean-field approaches.

We presented a minimal model of a population, including only two complementary phenotypic attributes. The model can be extended in a number of directions without making it computationally intractable. It would be easy to model different senses, along with different resource types that are easier to detect with one sense than another (for instance olfaction and vision). We expect that multiple phenotypes could naturally co-exist in such environments. It is unclear, when an environment would support generalists feeding on all resource types, or non-interacting, specialist species. We also assumed that resources are available until consumed. Resources with a finite shelf life are effectively scarcer, and agents would have to detect them more rapidly (consider an environment with few highly energetic resources that are only available for a short time). In this case the energy flux into the population is determined by the foraging strategy, and does not equal the total flux of energy into the system. Our model can also be extended to allow for communication between conspecifics, such as the waggle dance exhibited by honeybees [Seeley, 1995; Frisch, 1993]. The effectiveness of such communication could then have a strong impact on which strategies are selected [Kim et al., 2023].

Our model does not assume that the population optimizes a fitness function. Stochastic models, including agent-based models and birth-death processes, have been used to predict the impact of intra- and interspecific competition on the evolution of foraging strategies [Doebeli et al., 2017; Liang and Brinkman, 2022]. While deterministic models can also be used to predict how a population will evolve under intra- and interspecific competition [Chesson, 2000], they necessitate modeling the impact of competition, which is often mathematically intractable. However, stochastic models show that evolution does not necessarily lead to monotonic increases in fitness [Nowak and Sigmund, 2004], as they can display periodicity [Rubin et al., 2021] and the coexistence of multiple competing phenotypes [Rubin et al., 2023].

## Appendix

In §3.3 and §3.4, we employed Monte Carlo simulations to approximate equilibrium phenotype distributions in our agent-based model, which operates as a stochastic, Markovian process in a high-dimensional space 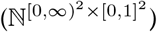 . We use around 10^2^ realizations to represent this process, which has a single absorbing state where all agents are extinct. However, with a sufficiently large energy influx, population size is stable over many generations, and we observe a quasi-stationary distribution of foraging strategies. Simulations indicate convergence to this distribution for different total population sizes, and resource densities.

Our model parameters are as follows: The side length of the domain, Ω, is unity. The persistent length *η* = 2^*−*4^. Initially, 10^4^ agents are sampled from a given phenotype domain ((*s*_*i*_, *a*_*i*_) *∈* (0, 0.2)^2^ in §3.4) and are located uniformly over the domain, Ω, and assigned initial energy *E*_*i*_(0) = 10. The coefficients of the rates of metabolic expenditures (2.1) are *c*_*s*_ = 1.6 10^2^ and *c*_*a*_ = 4. The coefficients of birth and death rates are *β* = 1, *δ* = 1, *δ*_0_ = 5 10^*−*3^, *α* = 4, *K*_*b*_ = 10, and *K*_*d*_ = 1. Offspring, when born, are placed at an initial point sampled from a normal distribution centered at their parents location, and with standard deviation *σ*_*x*_ = *σ*_*y*_ = 2.5 10^*−*3^. We non-dimensionalize the units as follows: the length is scaled by the side length of the domain, time is scaled by the birth rate *β*, and the energy is scaled by the threshold for the death rate *K*_*d*_. In simulations we used increments in time of size Δ*t* = 0.1.

We choose various mutation rates and environmental parameters in different simulations. In figure 2, we set *β* = *δ* = *δ*_0_ = 0 for the individual foraging process and the number of resources are *r* = 100, 200, 300. In figure 3, we set (*λ, F*) = (200, 1) with zero variability in attributes *σ*_*s*_ = *σ*_*a*_ = 0. We used Λ = *λF* = 200 for the mean-field approximation. *E*^*∗*^ can be determined using the parameters given above. In figure 4, we set 7 fixed metabolic cost values, *c* = 2^*−*6^, 2^*−*5^, *· · ·*, 2^0^ and 3 sets of environmental parameters (*λ, F*) = (12.5, 16), (100, 2), (400, .5) with the same energy influx to the system Λ = 200. There is no variability in attributes *σ*_*s*_ = *σ*_*a*_ = 0. We sampled 10^4^ phenotypes from the contour{*ϕ* : *C*(*ϕ*) = *c*}and repeated the simulation 10^2^ times until *T* = 2000. In figure 5, we used three sets of environmental parameters (*λ, F*) = (50, 4), (200, 1), (800, .25) for 20000, 500, 500 generations, respectively, and repeated simulations *M* = 10^2^ times. We set the variability to *σ*_*s*_ = *σ*_*a*_ = 3.75 *·* 10^*−*4^.

## Footnotes

1 We slightly abuse the term “equilibrium”, as this is actually a quasi-equilibrium. A domain with no agents is an absorbing state, and thus *N* = 0 is the only actual equilibrium. However, in the examples we describe this equilibrium was never attained in simulations, and is not relevant for our discussions.

