## Supplementary Materials for "Emergence of multiple foraging strategies under competition"

### S.1 Divergence of $L_0^2(r^*)$ with vanishing acuity

When the persistent length is larger than $L_0(r^*)$, the agent is likely to detect a resource before changing its search angle if it moves in the direction whose angle is in the arc formed by the agent at the tip, and the two tangents drawn from the agent's position to the circular disc of radius $a$ around the closest resource. The probability that upon turning an agent heads in such a direction is

$$P(a|r^*) = \frac{1}{\pi}\mathbb{E}\left[\arcsin \frac{a}{\min_{j=1,\cdots,r} d(\mathbf{x}_0, \mathbf{x}_j)}\right].$$

Using Jensen's inequality, we have the following lower bound

$$P(a|r^*) \geq \frac{1}{\pi}\arcsin\frac{a}{L_0(r^*)} \equiv \underline{P}(a|r^*),$$

which also gives the following bound for the average length of trajectories to a target

$$L(a|r^*) \leq \frac{L_0(r^*)}{P(a|r^*)} \leq \frac{L_0(r^*)}{\underline{P}(a|r^*)}.$$

Therefore, we have

$$\lim_{a\to 0} aL(a|r^*) \leq \pi L_0^2(r^*). \tag{S.1}$$

### S.2 Proof of Theorem 1

First, we find the critical points within the phenotype domain. In this domain, given our assumption about $g$, there exist two such points. Next, we will establish that only one of these points satisfies the conditions we assumed about $g$, which shows that this is the sole local minimum within the domain's interior, $\phi_{\text{opt}}$. Finally, we show there is only one local maximum at the boundary, $\phi_0$.

We first find local maxima (in fact only one maximum) in interior, say $(s, a) \in (0,\infty)^2$. Let $g(a|r) = 1/L(a|r)$, so that

$$\frac{\partial\varepsilon}{\partial s} = Fg(a|r) - 2c_s s, \quad \frac{\partial\varepsilon}{\partial a} = sFg'(a|r) - c_a.$$

Setting both derivatives to zero and solving for $s$ and $a$ gives,

$$s = \frac{F}{2c_s}g(a|r), \quad \frac{d}{da}\left[g^2(a|r)\right] = \frac{c_s c_a}{F^2}. \tag{S.2}$$

Let $s_{\text{opt}} = Fg(a_{\text{opt}}|r)/(2c_s)$. Taking the second derivatives of $\varepsilon$

$$\frac{\partial^2\varepsilon}{\partial s^2} = -2c_s, \quad \frac{\partial^2\varepsilon}{\partial s\partial a} = Fg'(a|r), \quad \frac{\partial^2\varepsilon}{\partial a^2} = sFg''(a|r).$$

and evaluating at the phenotype $\phi_{\text{opt}} = (s_{\text{opt}}, a_{\text{opt}})$ gives

$$\left.\frac{\partial^2\varepsilon}{\partial s^2}\right|_{\phi=\phi_{\text{opt}}} < 0, \tag{S.3}$$

and

$$\left.\frac{\partial^2\varepsilon}{\partial s^2}\frac{\partial^2\varepsilon}{\partial a^2} - \left(\frac{\partial^2\varepsilon}{\partial s\partial a}\right)^2\right|_{\phi=\phi_{\text{opt}}} = -\frac{F^2}{2}\frac{d^2}{da^2}[g(a_{\text{opt}}|r)]^2 > 0. \tag{S.4}$$

Eqs. (S.3) and (S.4) imply that $\varepsilon$ has a local maximum at $\phi_{\text{opt}}$.

We then find other maxima (and in fact it is also only one) on the boundary, $s = 0$ or $a = 0$. If $s = 0$, then

$$\varepsilon(0, a) = -c_a a, \tag{S.5}$$

which is maximized at $a = 0$. If $a = 0$, then

$$\varepsilon(s, 0) = -c_s s^2 \tag{S.6}$$

according to the fact that $\lim_{a\to 0} L(a|r) = \infty$. Again, it is maximized at $s = 0$. Therefore, there is only one candidate for the local maximum on the boundary, $\phi_0 = (0, 0)$. To show that $\phi_0$ is locally maximum, we prove that the directional derivative at $\phi_0$ (to the positive domain) is non-positive

$$\langle \mathbf{e}, \nabla\varepsilon|_{\phi=\phi_0}\rangle = -e_a c_a \leq 0, \tag{S.7}$$

where $\mathbf{e} = (e_s, e_a)$ for $e_{s,a} \geq 0$ and $e_s^2 + e_a^2 = 1$.

### S.3 Amount of resources at equilibrium given a homogeneous population

We can estimate the number of available resources at equilibrium, $r^*$, when agents have high acuity or there are many resources in the domain so that the agents can detect resources immediately, $a \gg L_0(r^*)$. Since we expect that equation (3.2) holds in this parameter regime, substituting equation (3.2) into equation (3.6) and rearranging gives

$$L_0(r^*) = \frac{s \cdot n^*(\phi)}{\lambda}. \tag{S.8}$$

The right-hand side is determined by phenotypic and environmental parameters. Since the closed form of $L_0(r^*)$ is known (see the electronic supplementary material),the number of resources at equilibrium $r^*$, can be estimated by solving (S.8) for $r^*$. Since $L_0(r)$ is a decreasing function, $r^*$ increases as the value of the right-hand side decreases. As expected, many resources are available at equilibrium if resources are generated rapidly, agents move slowly, or there are few agents in the domain.

### S.4 Expression for $L_0(r)$

Since our domain is periodic and thus translation invariant, it suffices to calculate

$$L_0(r) = \mathbb{E}\left[\min_{j=1,\cdots,r} d(0, \mathbf{x}_j)\right], \quad \mathbf{x}_j \in [0,1]^2$$

Introducing the random variables

$$\mathcal{D}_r = \min_{j=1,\cdots,r} d_j, \quad d_j = d(0, \mathbf{x}_j), \quad \mathbf{x}_j \sim U[0,1]^2,$$

for $j = 1, \cdots, r$, the cumulative density function of a distance between an agent and a resource satisfies

$$\begin{aligned} 1 - H(y) &:= \mathbb{P}[d_j \leq y] \\ &= \begin{cases} \pi y^2, & y \leq 1/2 \\ \sqrt{4y^2 - 1} + 4y^2\theta(y), & 1/2 < y \leq \sqrt{2}/2 \end{cases} \end{aligned} \tag{S.9}$$

where

$$\theta(y) = \frac{\pi}{4} - \arcsin\frac{\sqrt{4y^2-1}}{2y}.$$

Since $d_j$ are independent and identically distributed, we have

$$\begin{aligned} \mathbb{P}[\mathcal{D}_r > y] &= \mathbb{P}[d_j > y, \forall j = 1, \cdots r] \\ &= \prod_{j=1}^{r} \mathbb{P}[d_j > y] \\ &= H^r(y). \end{aligned}$$

Then $L_0(r)$ can be written by

$$L_0(r) = \int_0^{\sqrt{2}/2} y \frac{d}{dy} \left(1 - H^r(y)\right) dy$$

Integration by parts gives

$$L_0(r) = \int_0^{\sqrt{2}/2} H^r(y) dy. \tag{S.10}$$

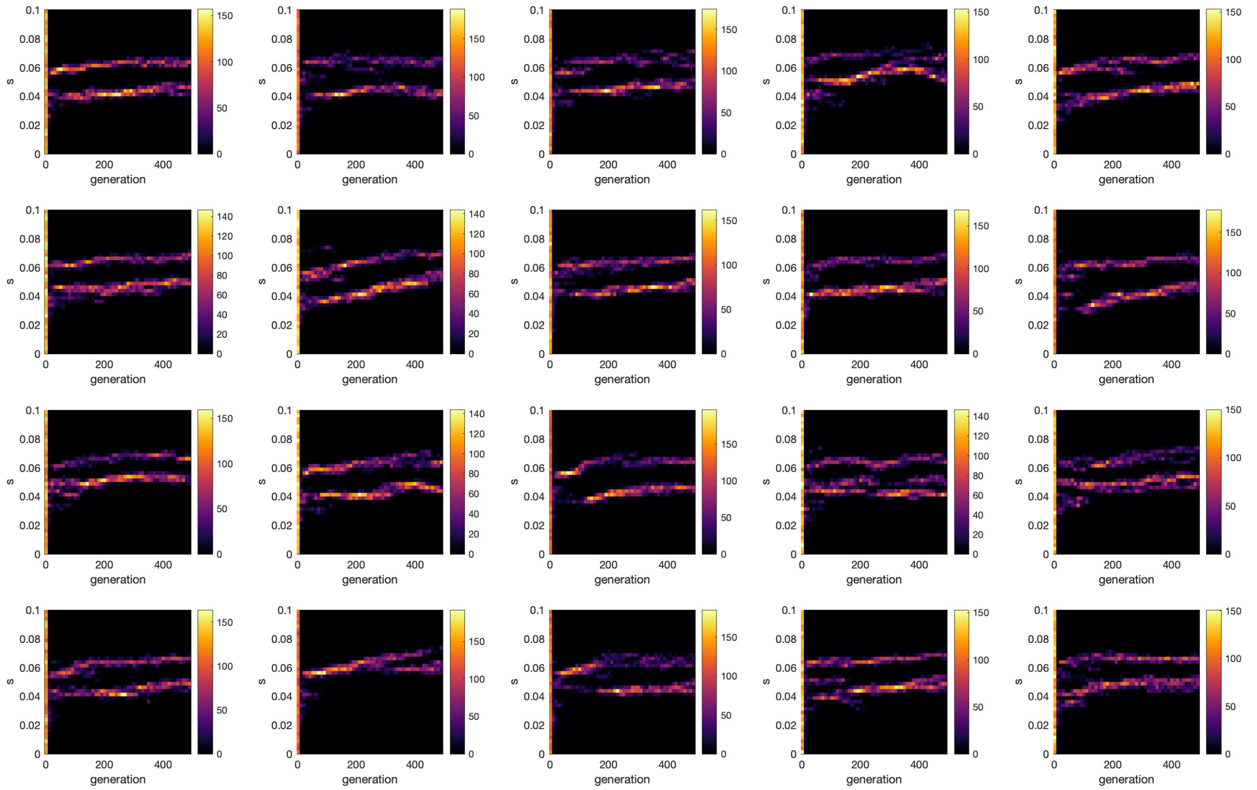


**Figure S1.** Evolution of foraging strategies in the moderate environment ($F = 1$, $\lambda = 200$). Each panel represents a single Monte Carlo simulation of evolution of foraging strategies and shows a histogram of foraging speed over 500 generations. Almost all panels show persistent coexistence of distinct foraging speeds (and thus foraging strategies) over generations. Similar results hold for acuities.

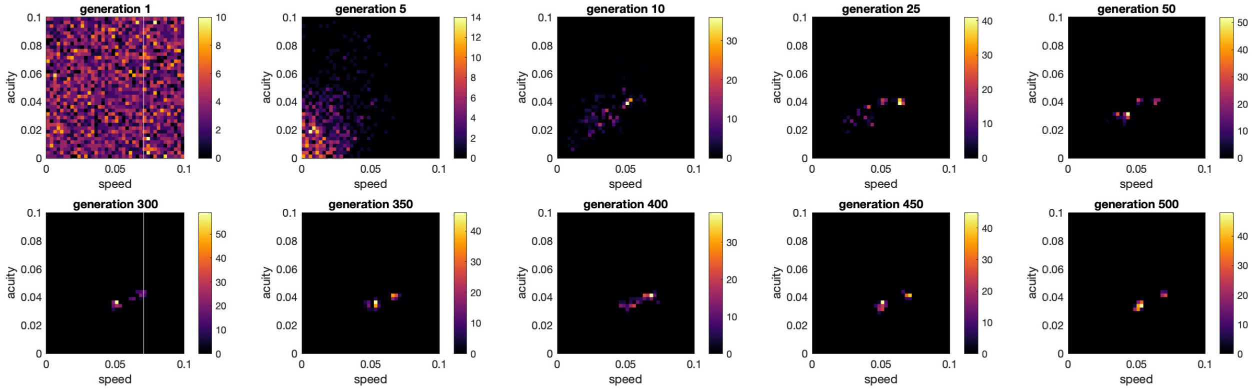


**Figure S2.** Typical evolution of foraging strategies in the moderate environment ($F = 1$, $\lambda = 200$). Each panel shows a phenotype histogram for a fixed generation. This shows that distinct foraging strategies coexist over many generations in the moderate environment.

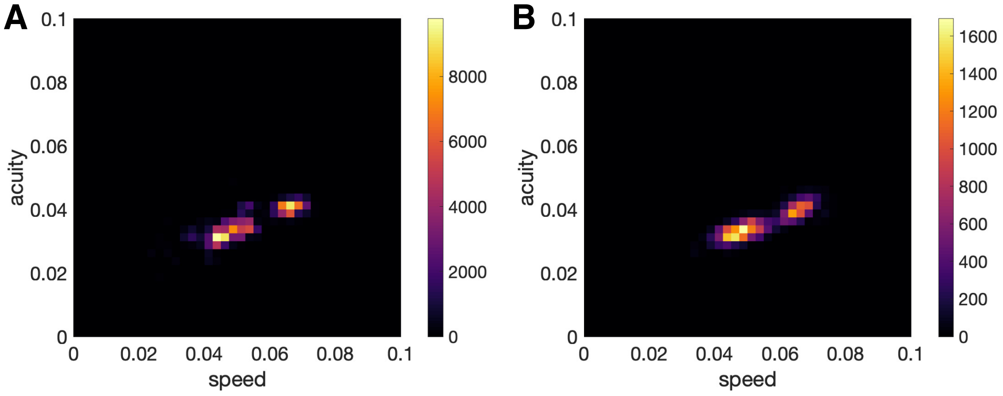


**Figure S3.** Ergodicity of simulations in the moderate environment ($F = 1$, $\lambda = 200$). (A) Averaged histogram of foraging strategies over time. The histogram is generated by pooling over generations. (B) Averaged histogram of foraging strategies over event space. The histogram is generated by pooling over many Monte Carlo simulations at a given time after the start of the simulation.

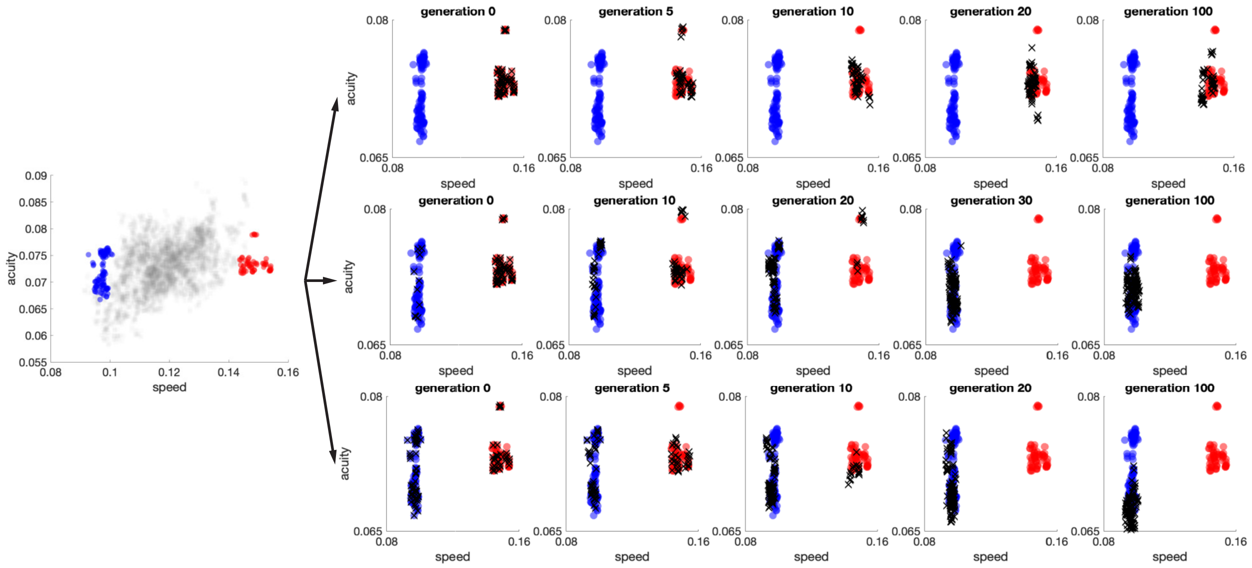


**Figure S4.** First, from many simulations in low $\lambda$ and high $F$ environment (*gray squares*), we select two extreme simulations with high metabolism (*red circles*) and low metabolism (*blue circles*). We start the simulations with three initial conditions from those two extremes: All phenotypes from the high metabolism distribution (*top panels*); equally distributed phenotypes from the two distributions (*bottom panels*); phenotypes mostly from the high metabolism distribution (*middle panels*). Over generations the population converges to a low metabolism state even when the initial number of this phenotype is low. This shows that the the less costly phenotype outcompetes the more costly one in this environment in direct competition. However, since the population size is low, it takes an extremely long time for a population starting in the high state to converge to a low state.

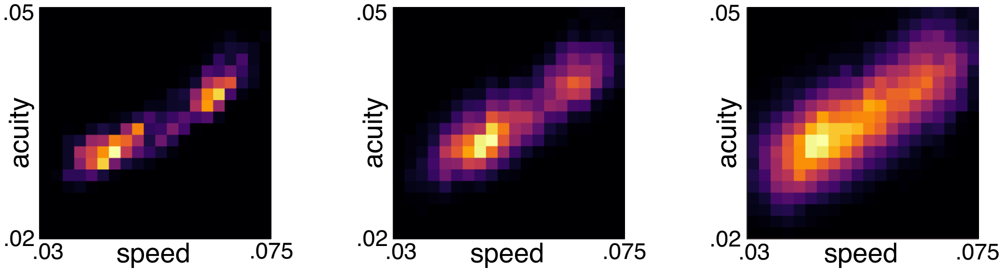


**Figure S5. Phenotype distribution depends on the variability in the offspring phenotype.** As the variability increases (*from left to right panels*), the two modes of the phenotype distribution at equilibrium vanishes into the single mode distribution. We set the environmental parameters $\lambda = 200, F = 1$ and changed the variability of offspring by using the three values $\sigma_s = \sigma_a \equiv \sigma = 1.25 \cdot 10^{-4}, 4.375 \cdot 10^{-4}, 10^{-3}$ in simulations.
